# Scalability of large neural network simulations via activity tracking with time asynchrony and procedural connectivity

**DOI:** 10.1101/2021.06.12.448096

**Authors:** Cyrille Mascart, Gilles Scarella, Patricia Reynaud-Bouret, Alexandre Muzy

## Abstract

We present a new algorithm to efficiently simulate random models of large neural networks satisfying the property of time asynchrony. The model parameters (average firing rate, number of neurons, synaptic connection probability, and postsynaptic duration) are of the order of magnitude of a small mammalian brain, or of human brain areas. Through the use of activity tracking and procedural connectivity (dynamical regeneration of synapses), both computational and memory complexities of this algorithm are proved to be theoretically linear with the number of neurons. These results are experimentally validated by sequential simulations of millions of neurons and billions of synapses running in few minutes using a single thread of an equivalent desktop computer.

## 1 Introduction

There are more and more vast research projects, whose objective is to simulate brain areas, or even complete brains, in order to better understand their functioning. Examples are the Human Brain Project (HBP) in Europe, the Brain Mapping by Integrated Neurotechnologies for Disease Studies (Brain/MINDS) in Japan or the Brain Initiative in the United States. Several approaches can be considered for this purpose. There is the biochemical approach (see for example [38]), which cannot succeed for systems as complex as the brain. A more biophysical approach has been used (see for example [13]) where cortical barrels have been successfully simulated. This approach is limited to about 10^5^ neurons. However, the human brain contains about 10^11^ neurons while a small monkey, like marmosets [6], already has 6 · 10^8^ neurons [22] and a larger monkey, like a macaque, has 6 · 10^9^ neurons [22].

To simulate such huge networks, the models have to be simplified. In particular, a neuron has no physical form and is just represented by a point in a network, possibly with a certain voltage. The Hodgkin-Huxley equations [33] are able to reproduce the shape of the action potential by taking into account the dynamics of the ion channels, but the complexity of these coupled equations [18], makes the system quite difficult to simulate for huge networks. If the dynamics of the ion channels are neglected, the simplest voltage model is the Integrate-and-Fire (IF) model. With such models, it has been possible to simulate on supercomputers a neural network reaching about 68 · 10^9^ neurons [55].

However, there is another range of simplified models, which can simulate such massive networks. Indeed, we can use much more random models to reproduce the essential dynamics of neurons: their firing pattern. Randomizing not only the connectivity graph but also the dynamics on the graph makes the model closer to real data and explains to some extent their variability. The introduction of randomness is not new and has been done in the models mentioned above: Hodgkin-Huxley [12] or Leaky Integrate-and-fire (LIF) [29].

Here we want to focus on particular random models: point processes [51], which model only spike trains. Point processes have usually the property of time asynchrony, *i.e.*, two different neurons cannot produce spikes at exactly the same time. This includes Hawkes models and its variants such as generalized linear models, Wold processes, Galves-Löcherbach models, and even some random LIF models with random or soft thresholds [51, 4, 44, 41, 42, 14, 46]. As one can see in these references, all these point process models have been used to study different real datasets. Some of these point process models can be related to IF models. Indeed, as noted in [15] (Chapter 9), IF models can be made noisy, using “escape noise”. In an escape noise model, IF neurons can fire even if the threshold has not been reached or can remain quiescent even if the threshold has been transiently exceeded. As indicated in the same book (chapter 10), this formulation is very close to the generalized linear models [41], which are in fact known in mathematics as Galves-Löcherbach models [14] or the more classical version without reset: Hawkes models [44]. In this sense, Hawkes processes, which are used in the present work, can be seen as a noisy version of an IF model.

The time asynchrony property of point processes [2, 4], combined with the sparsity of the graph led to a new algorithm for point process network simulation, whose computational complexity is theoretically controlled. We called this algorithm ATiTA, for Actitivity Tracking with Time Asynchrony, in [32]. Thanks to time asynchrony and activity tracking [34], we showed in particular that, if the graph is sparse, the complexity cost of the calculation of a new point in the system is linear with the number of neurons. However the memory cost of this former work was too high to reach networks of 10^8^ neurons.

In a preliminary work on the mathematical aspects of the mean field limits of LIFs [17], we also found a way to optimize the memory cost in the framework of random graphs: it is enough not to keep in memory the whole network but just to regenerate it when needed. However, this trick was not put into practice and its memory cost was not theoretically calculated in [17]. Both new aspects are treated in the present article. This approach is inspired by dynamic-structure discrete event systems [36]. It has already been applied to track the activity and addition/deletion of cells and neighborhood connectivity in cellular automata [35]. Recently, the same idea under the name of procedural connectivity, has been successfully applied on LIF models using Graphics Processing Units (GPU) [26]. Thanks to GPU parallel programming, and without using time asynchrony, the authors of [26] were able to simulate in parallel a network of order 10^6^ neurons and 10^10^ synapses in a few tens of minutes on a GPU graphics card running thousands of parallel threads.

In comparison to [26], by combining procedural connectivity with activity tracking and time asynchrony, the new algorithm of the present work, called ATiTAP, obtains a smaller memory cost and a range of computational costs, which are much smaller than the sum of the costs of thousands of threads, and this, by using a single thread.

Indeed, classical parallel computing generally uses a discrete simulation time and computes for all neurons (or synapses) what happens at each time step in a parallel way. However, the spikes must be transmitted between pre and post synaptic neurons, between two time steps. With more advanced techniques such as hybrid parallel computation [19], the spikes are computed in parallel for each neuron thanks to discrete event programming, which allows a large precision, while the exchange at the level of the synapses can be done at a less fine scale in a discretized way [37, 3, 45, 50]. Using hybrid parallel computations, in [24] the authors were able to simulate a network of order 10^6^ neurons and 10^10^ synapses with parallel supercomputers capable to run in parallel tens of millions of threads. Nevertheless, in both cases (classical or hybrid parallel computation), the synchronization between neurons is necessary to propagate the spikes at the synapses, even if might be done at a rougher time scale than the precision for the hybrid algorithm.

With ATiTAP, as a result of time asynchrony, we can exploit discrete event computations [56, 34] to track the activity, not of each neuron independently, but of the whole neuronal network, over time, by jumps (from one spike in the network to another spike in the network). Synaptic transmissions are updated on the fly only for the neurons impacted by a spike. Therefore, a single thread can be used to sequentially compute each spike and each synaptic transmission. Being alone, the single thread can then compute the dynamics of the entire network as quickly as possible without waiting for the synchronization with other threads. This reduces the overall computation time (compared to the summed time of all threads), which makes our sequential computation technique competitive with parallel computation techniques. Indeed, being able to simulate an entire large neural network using a single thread opens the possibility of using the other threads to run simulation replicas in parallel. It is then possible to simulate many models in parallel instead of a single model in parallel, with reasonable execution time.

To summarize, ATiTAP, the new algorithm proposed here, allows simulating large neural networks through activity tracking with time asynchrony and procedural connectivity. Thus, a realistic network of 10^8^ neurons can be sequentially simulated on a single thread, with memory and computational costs roughly comparable to other available algorithms (which require a massive parallel implementation on multiple threads) [24, 26, 47]. Moreover, these costs can be controlled analytically in advance: in particular, they remain linear with the number of neurons in the network.

In Section 2, after discussing more precisely the differences and links between Hawkes processes and classical Integrate-and-Fire, we explain the main features of the benchmark (the hybrid simulation algorithm for IF [19]) and the ones of our new algorithm ATiTAP. We also derive the theoretical memory and computational costs and compare it to simulations on a realistic range of parameters with respect to human brain areas. In Section 3, we give the main mathematical ingredients to calibrate our simulations. In Section 4, we discuss the results we obtained in view of the various benchmarks of the existing algorithms to simulate huge realistic neuronal networks.

## 2 Results

### 2.1 Models features

As said in the introduction, some point processes and especially Hawkes processes have some common features with more classical Integrate-and-fire models. Point processes are usually defined by their conditional intensities, which describe the probability for a spike to happen on a given neuron, given every other spikes that have happened in the network in the past [4]. It has been shown (see for instance [23] with experiments on motor neurons), that the conditional intensity of a spike train is a function of the membrane voltage. From a model point of view, we can assimilate more or less one to the other: the higher the voltage, the more likely the neuron spikes. This is also the spirit of the equivalence derived in chapters 9 and 10 of [15]. In this sense, one derives an equivalence between the formula of the conditional intensity of a Hawkes process and the voltage dynamics of an Integrate-and-Fire model. Modeling the activity of *M* neurons by a Hawkes process means that the conditional intensity of neuron *i*, which models the spike train *N^i^*, is given by

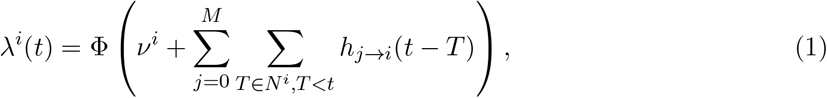

In this formula, from a point process point of view, *ν_i_* represents the spontaneous activity of the neuron *i* if the other neurons do not fire, whereas *h_j→i_* is the interaction function, that is *h_j→i_*(*u*) is the increase (if positive) or decrease (if negative) that the firing rate of neuron *i* suffers due to a spike on *j*, which has taken place *u* seconds before. The increasing function Φ is the transfer function, which is usually the positive part, *i.e.* Φ(*x*) = max(*x*, 0). Indeed, if the interaction functions *h_j→i_* are too negative, the overall sum might be negative and the positive part ensures that the conditional intensity is just null in this case: this corresponds to the fact that the neuron is so inhibited that it does not emit any spike anymore. Some authors also uses the exponential transform Φ(*x*) = exp(*x*) (see [41] for instance).

From a voltage/IF point of view, we can interpret

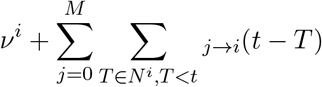

as the voltage membrane of the neuron, *ν^i^* being the resting potential and the functions *h_j→i_*, the respective excitatory or inhibitory post-synaptic potentials that are summed after each received spikes. In the IF model, especially the one used in [19], these functions are solutions of differential equations (the most classic, the exponential, being represented in blue on Figure 1, less classic functions with synaptic delays for the transmission being represented in black).

**Figure 1:**
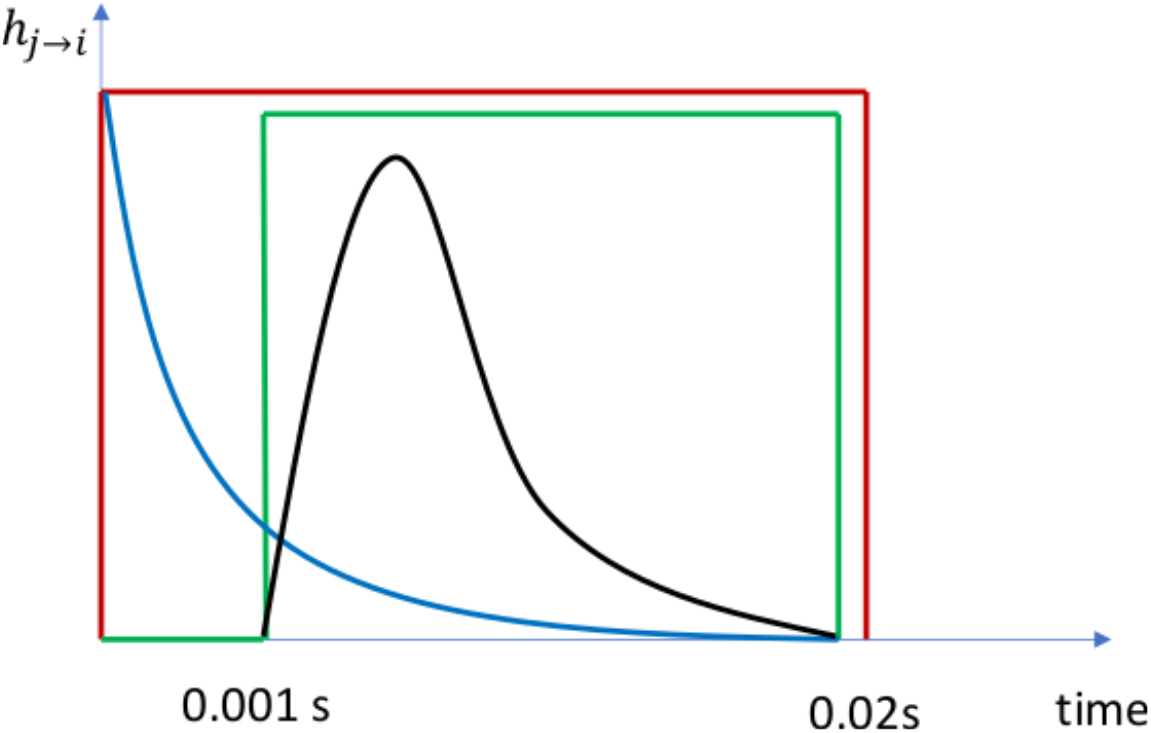
Examples of interaction functions *h_j→i_* from a neuron *j* to a neuron *i:* in red a piecewise constant function with no delay; in green a piecewise constant function modeling a synaptic transmission delay; in black a continuous interaction function with the same synaptic delay; in blue the exponential interaction function, obtained in the most classic Integrate-and-Fire models.

Once this parallel is done, there are several features that Hawkes processes have that might help to understand why we choose this model over various other representations.

First of all, because Hawkes processes are defined by a conditional intensity, they are simple [2] and therefore do possess the time asynchrony property, that is, two neurons cannot produce spikes at exactly the same time. This property is not satisfied by classical Integrate-and-Fire models, which fire each time the voltage of a given neuron exceeds a given threshold. It has been shown mathematically that unrealistic blow-up phenomena can appear for IF because of this absolute rule: a massive proportion of the neurons can exceed the threshold at the exact same time, whereas Hawkes models do not have this caveat [5, 7].

Second, and this is true for all point processes defined by a conditional intensity, one can always thin a rougher process into a more complex one [39, 32]. Indeed as soon as we are able to simulate a process with a larger intensity, we can reject some of the points, via a particular mathematical procedure called *thinning*, to create a process with a smaller intensity.

Third, in the case of excitatory interaction functions and transfer function Φ(*x*) = *x*, we can easily predict mathematically the number of spikes that will be produced (see Section 3 for more details). This helps us design beforehand numerical experiments with a given highly variable distribution of the firing rates, as one can observe on real data.

Finally, in the case of non negative piecewise interaction functions, thinning is not necessary [32] and we are able to precisely compute the theoretical memory and computational costs (see Sections 2.3 and 2.4) as a function of *e.g. M*, the number of neurons; *d* the average degree of the network; 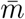, the average firing rate in the network; *τ*, the largest possible interaction duration, that is the maximal support of the functions *h_j→i_* and *A*, the maximal number of breakpoints in the piecewise constant functions *h_j→i_*.

In the sequel, we restrict ourselves to non negative piecewise constant interaction functions (and Φ(*x*) = *x*) for the computation and simulation. Note that taking negative piecewise constant functions is not a problem for ATiTA and ATiTAP, and that the resulting process will always have less points than the one without inhibition. Memory cost and numerical complexities are therefore upper bounded by the pure excitatory case, where negative interaction functions are discarded. Also it is always possible to thin this process into a more complex one with smoother interaction functions for instance, as long as the interaction functions are upper bounded by piecewise constant functions (see Figure 1). Again this procedure will only diminish the number of points. However the theoretical complexity of this thinning step cannot be well evaluated. Therefore the benchmarks that are given in the present work have to be understood as a proof of concept that such huge simulations can be done using a single thread. Of course, we can also make the interaction functions smoother by increasing the number of breakpoints of the piecewise interaction functions (*A*). However, it is likely that this would be more time consuming than pure thinning.

### 2.2 Description of the algorithms

#### Parallel simulation and the hybrid algorithm

The simulation of large neural networks is usually performed in parallel with regular synchronizations of all the computing cores. Of course, this depends on both the mathematical model and the simulation algorithm. However, for most models, differential equations are used to compute the time evolution of the membrane voltage for each individual neuron. Usually, these equations are (approximately) solved by classic discrete-time numerical schemes [26]. One of the most advanced algorithms nowadays to simulate neuronal networks is called hybrid parallel computing [19] and it works with an Integrate-and-Fire model. It supersedes the previous discrete-time numerical schemes by using a discrete-event simulation to directly produce the next spike of a neuron once all synaptic transmissions to this neuron have been planned, and this with large precision.

The hybrid parallel algorithm is represented on the top part of Figure 2 (a. and b.). There are two ways to understand the computations: either from a neuron point of view or from a hardware point of view. From a hardware point of view we refer to Processing Units (PUs) as the minimal hardware processing unit where one or more threads are executed, which would be a single Central Processing Unit (CPU) core or an Nvidia GPU Streaming Multiprocessor (SM). From a neuron point of view, when a presynaptic neuron emits a spike at continuous^1^ time *t* (red ticks on Figure 2a), the postsynaptic neurons receive this information at time *t* + *T_com_* (orange dots on Figure 2a), where *T_com_* is the minimal synaptic transmission delay in the network. Between two synaptic transmissions, the membrane potential of a neuron evolves independently from the other neurons and can be calculated in parallel (green dashed lines on the figure 2a). From a PU point of view (*cf*. Figure 2b), a PU takes in charge the parallel computations of a group of neurons. Because of the minimal delay *T_com_*, every multiple of *T_com_*, the PUs must synchronize to be able to anticipate, over the next interval of length *T_com_*, the synaptic transmissions that are going to take place, before computing independently the membrane voltage. Notice therefore that since the firing rate is quite low in the network, a lot of PUs could have continued their computations without synchronization, but are forced to wait. Hence the load of each PU is sub-optimal on average.

**Figure 2:**
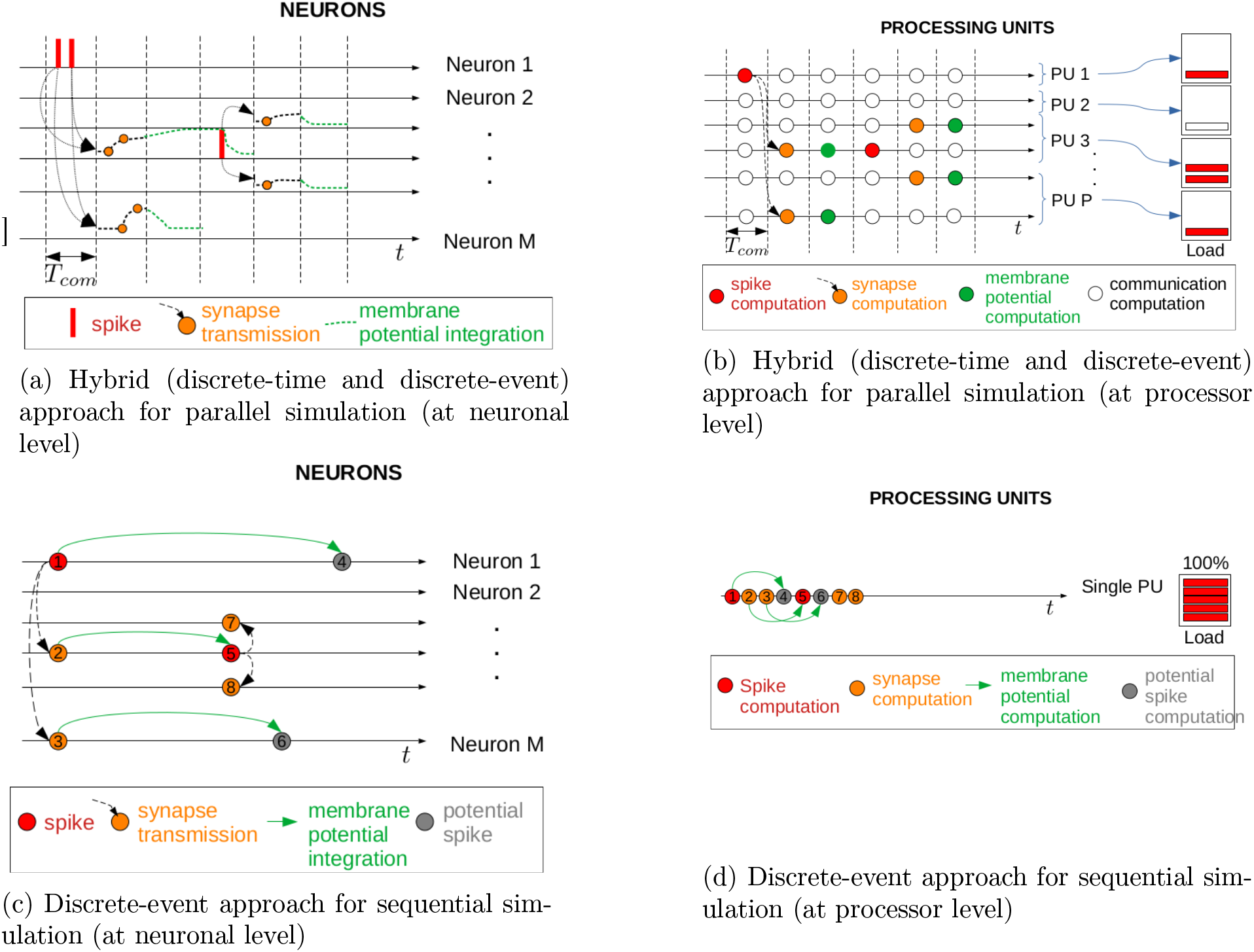
Schematic view of the hybrid algorithm [19] (Figures (a) and (b)) and ATiTA(P) (Figures (c) and (d)). Figures (a) and (c) represent the neuronal network point of view, whereas the PU point of view is displayed in Figures (b) and (d). Everywhere, spikes are in red, synaptic transmission events in orange, prediction of next spike in green, thanks to integration of membrane potential or intensity. For ATiTA(P), also grey refers to potential spikes that are computed and then discarded. In (a) thanks to the synaptic delay of size *T_com_* spikes are received by the neurons in the next bin of size *T_com_* and then integrated to compute the membrane potential. In (b) the computations per PU are then done per bin of size *T_com_* and need to be synchronized every *T_com_*. Depending on the number of PUs, it is likely that some PUs wait the other ones without computing much on each thread and therefore their load is low. In (c), for ATiTA(P), the discrete-event approach is used at the network level: the computation makes jumps to the next potential spikes. The smallest one is kept as the actual next spike. Then synaptic transmission, update of the corresponding intensities and new computations for the next potential spikes, are done only for the post synaptic neurons. In (d), the different operations of (c) are put in the order of the successive operations that are done by the single thread on the single PU, which has therefore a full load over time. Note that both algorithms (hybrid and ATiTA(P)) have a time precision, which can be the classical numerical precision of 10^15^ and in this sense, they both compute continuous times.

#### ATiTA and ATiTAP

In [32], we proposed a discrete-event algorithm, ATiTA, to simulate point processes with stochastic intensities. This algorithm is based on the theory of local independence graphs [9], which corresponds to the directed neuronal network in our present case. ATiTAP follows the same principle and we refer to both algorithms as ATiTA(P). Computationnally, ATiTA(P) work as follows (see Figure 2c).

From a neuronal network point of view, the spike events happen in continuous time in the system (up to the numerical precision). Once a spike on a particular presynaptic neuron happens (red dots in Figure 2c), the postsynaptic neurons are updated (orange dots in Figure 2c). The presynaptic and the post synaptic neurons compute their respective intensities and forecast their evolution (green arrows in Figure 2c) if nothing in between occurs in the system. They are therefore able to forecast their potential next spike (gray dots in Figure 2c). The algorithm maintains a scheduler containing all potential next spikes on all neurons. This particular data structure [32] can efficiently maintain a sorted list of numbers. ATiTA(P) decides that the next neuron to fire effectively is the one corresponding to the minimum of these potential next spikes. For more details, we refer to [32]. The computational gain comes from the fact that neurons that are not firing a lot, do not require a lot of computations either. In particular we do not have to update all neurons at each spike but only the pre and post synaptic neurons that are involved in the spiking event. This is the main difference with the hybrid algorithm detailed above, which requires each PU to wait for the others to finish at each bin of size *T_com_*, even if it does not receive any input. Note that the whole algorithm is possible only because two neurons in the network will not spike at the same time: the whole concept is based on time asynchrony to be able to jump from one spike in the system to the next spike in the system. Of course, this is true only up to numerical precision: if two potential next spikes (gray dots on Figure 2c) happen at the exact same time with double resolution 10^−15^, by convention the neuron with the smaller index is said to fire. But the probability of such event is so small that this is not putting the simulation in jeopardy. Also note that this does not prevent neurons to eventually synchronize quite frequently over few milliseconds, as defined for instance in [52] and the references therein.

From a PU point of view (*cf*. Figure 2d), a single thread of a single PU computes sequentially (potential) spikes, synaptic and membrane potentials for all the neurons in the network. Notice that the load of this single PU is always optimal, as it is computing all the activities in the network. At the difference with the hybrid algorithm, no load balancing is required to attribute threads and PUs to groups of neurons. Computational performances only depend on the capacity of a single PU to compute activities as fast as possible.

The additional feature of ATiTAP with respect to ATiTA is the procedural connectivity [17, 26]. One of the memory burden of the hybrid algorithm and of ATiTA, comes from the fact that a classic implementation stores the whole connectivity graph, which is huge for brain scale models. If the connectivity is the result of a random graph and that each presynaptic neuron is randomly connected to its postsynaptic neurons, one can store the random seed instead of the result of the random attribution. Hence the whole graph is never stored in full but only regenerated when need be. The random connectivity is regenerated at each spike taking advantage of the deterministic nature of the pseudo-random generator used in the simulation. Storing the generator initial seed, the seed of each neuron is computed based on initial seed value and neuron index (see Figure 3). With this method, only the initial seed is stored in memory. Of course this dynamic regeneration at each spike has a cost in terms of time complexity, but this cost is negligible with respect to the other computations that need to be made and this saves memory. In the sequel, the random graphs that are considered are Erdös-Renyii (see Section 3).

**Figure 3:**
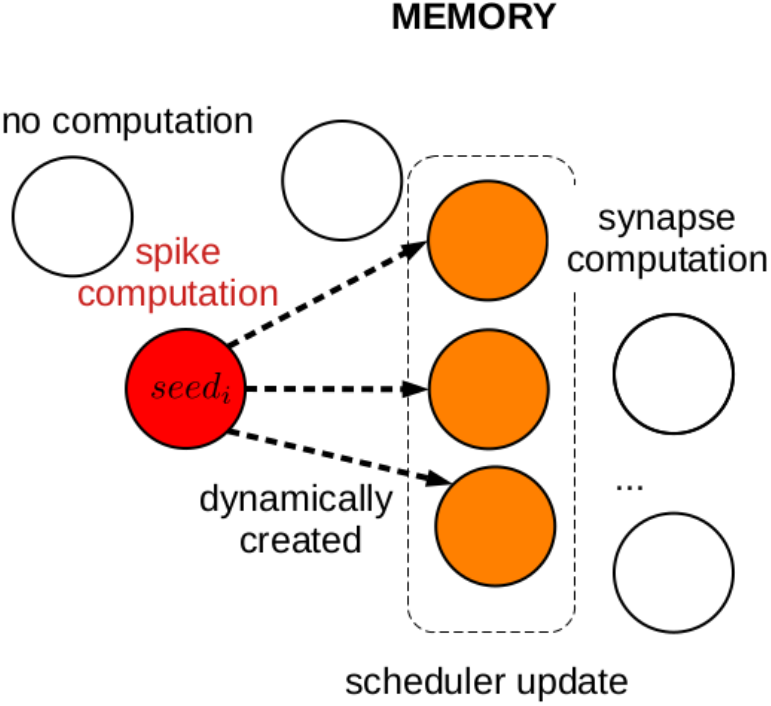
Time asynchrony and procedural connectivity at both computational and memory levels. When a spike occurs in neuron *i*, corresponding seed is computed based on neuron index *i* and the indices of the corresponding post-synaptic neurons are then generated. Later, the next potential spikes of these post-synaptic neurons are computed. Colors represent the computational order of magnitude: white meaning no computations, orange corresponding to the computation of the next potential spike and red to the computation of the next potential spike plus the seed computation and local regeneration of the graph.

### 2.3 Theoretical computational cost of ATiTA(P)

In [32], an accurate estimate of the complexity of ATiTA has been derived (*cf*. Equation 7 of [32]). This can be completed to compute the computational complexity of ATiTAP, when the procedural connectivity step is added. The overall time complexity of our algorithm is of the order of

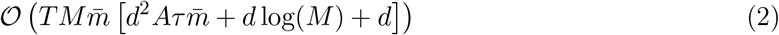

More precisely, since *M* is the number of neurons, 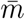 the average firing rate and *T* the simulation time, the factor 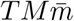 corresponds to the average number of points that are produced by the algorithm. For each of these points, one needs to decompose the algorithm in a few steps to understand the different elements of this complexity formula.

a. *Update of the intensities of the presynaptic neuron and of all the d (in average) post-synaptic neurons and computation of their possible next point:* The intensity of a given neuron is piecewise constant, more precisely it is a sum of (in average)*d* piecewise constant functions, corresponding to each interaction functions (see (1)). Moreover a term like ∑_*T∈N^j^,T<t*_ *h_j→i_*(*t–T*) is piecewise constant in *t* with as many breakpoints as: *A* (number of breakpoints in *h_j→i_*) times the number of spikes that are appearing in a range *τ* (support of the function *h_j→i_*). So globally, the intensity of a given neuron is a piecewise constant function with (in average) 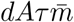 breakpoints. Updating *d* + 1 intensities and computing for each of them, their next potential point is therefore proportional to 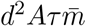.
b. *Update of the scheduler containing all the possible M next points*: Updating one time in a scheduler requires log(*M*) iterations. Hence updating *d* + 1 points in the scheduler requires *d* log(*M*) iterations.
c. *Procedural connectivity*: This step is the only difference between ATiTA and ATiTAP. It does not exist for ATiTA. For ATiTAP, at each spike, we regenerate the *d* (in average) postsynaptic neurons of the random graph. This costs *d*. Note that step (c) is negligible with respect to step (b) and therefore ATiTA and ATiTAP have roughly the same computational complexity.

Note that for general point processes, the main change in this evaluation would be the complexity of the intensity in step (a), which would also impact the computation of the next point. If thinning is involved, that is if we simulate a rougher process and then precisely reject all the unnecessary points, the global factor will be larger than the actual number of points 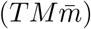, since it would count the number of points of the rougher process. However, if the evaluation of the more complex intensity at a given point is not too time consuming [39], the complexity (for a given point of the rougher process) of step (a), (b) and (c) can be roughly the same.

### 2.4 Theoretical memory cost of ATiTA(P)

In [32], the memory cost of ATiTA was not computed. Here we evaluate this cost for both ATiTA and ATiTAP. There are several costs, some necessary whatever the aim of the simulation, some depending on what is the purpose of the simulation.

a. *Library cost:* As in [27, 24], some libraries and other code need to be stored in memory. We denote this cost by *ρ*. Since there is only one PU and we are not using a parallel simulation, we pay this cost only once.
b. *Random seed:* For ATiTA or ATiTAP, random numbers are necessary to produce the points of the process. For reproducibility purpose, one can store the first random seed and then obtain the other ones based on neuron indexes. Hence only the first one needs to be stored. This cost is *η*.
c. *Connectivity cost:* For ATiTA, one can use the Yale format [11] to store the synaptic connections of the neural network. This format consists of storing for each pre-synaptic neuron the set of post-synaptic neurons (both of them represented as indices). So the memory cost if *dMω*, with *ω* the number of bytes to store an index (integer or long integer). For ATiTAP, we also need to store the first seed for the generation of the random graph. Then, the other ones, for each neuron of the graph, are obtained thanks to their index (see Figure 3). Finally, at each iteration of the algorithm we need to store, in average, the *d* indices of the *d* post synaptic neurons that are regenerated on the fly. The whole cost is therefore *η* + *dω*.
d. *Parameters cost:* In general, for ATiTA, it should be necessary to store all the *ν_i_*’s and all the *h_j→i_*’s that are non zero. This would cost roughly *MdAϵ*, where *ϵ* is the number of bytes necessary to represent a spiking time. In the set-up of the present simulation, for ATiTAP, the *ν_i_*’s are different to have a large heterogeneity in the spiking rates and we use the same interaction functions for all neurons as soon as they are non zero. The cost is therefore (*M* + *A*)*ϵ*.
e. *Storage of the intensities:* As said previously in Section 2.3, an intensity has in average 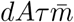 breakpoints. Hence, storing all the *M* intensities roughly costs 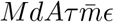.
f. *Scheduler of the possible next spikes:* This scheduler is of size *M*, hence the cost is *Mϵ*.
g. *Storage of all the generated points:* Depending on the problem, one might want to store all the spikes that have been generated, or only summary statistics (like firing rates, etc.) that are less costly. In the case where we want to store all the generated points, the cost is 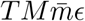.

Hence without counting case (g), we have for a general ATiTA algorithm with various interaction functions a memory cost of

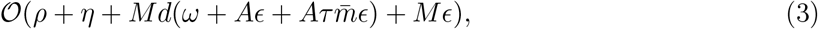

whereas ATiTAP with one common interaction function is of memory cost

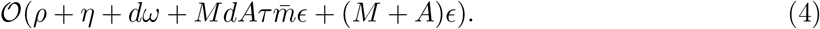

Usually, *η* = *ω* = *ϵ* = 64bits, the precision of a double / long integer type.

### 2.5 Choice of brain scale parameters

Because of the precision of actual measurements and their intrinsic variability, it is difficult to estimate quantitatively both physiological (number of synapses per neuron, etc.) and dynamic parameters (average firing rate, etc.) of neuronal networks in primates [22] and humans [21]. Only rough estimates are available. We give in this section, rough estimates of the main quantities of a human brain in order to see the scalability of ATiTA(P) to reach human brain areas.

To our knowledge, the best documented region of the human brain is the (neo)cortex. Based on the structural statistics (number of neurons and synaptic connections) of neuronal networks in the (neo)cortex, we extrapolate here their representative parameter values for the whole brain.

The *firing rate of a neuron* in the brain can be estimated by the limited resources at its disposal, especially glucose. Measures of ATP consumption have shown (see [30]) that the firing rate of a neuron in human neocortex can be estimated around 0.16*Hz*. Still based on ATP consumption, only 10% of the neurons in the neocortex can be active at the same time. So it seems coherent to choose an average of 0.16*Hz*. These values can be extrapolated to the whole brain^2^, as follows.

The neocortex represents 80% of the volume of the brain [49] and consumes 44% of its energy [30]. Considering that the energy consumed by the brain is proportional to the firing rate of the neurons, we obtain that

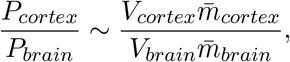

with 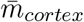 the mean firing rate of individual neurons in the neocortex (resp. in the brain) and *V_cortex_* the volume of the neocortex (resp. in the brain). The *average firing rate of the brain* then consists of 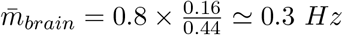 per neuron.

This average firing rate should not be confused with the fact that particular neurons may have a much larger firing rate. Particularly, groups of neurons synchronize together to achieve a particular cognitive task: this is the concept of neuronal assemblies [16]. In an assembly, neurons can usually increase their rates to tens *Hz* (possibly 50*Hz*) over a short duration. Therefore, we choose a firing rate distribution where most of the neurons have a firing rate of 0.3*Hz* but some have a much higher firing rate (up to 50*Hz*) using an heavy tailed distribution (see Section 3).

The average number of synaptic connections in a human brain is hard to estimate and depends heavily on the neuron types and brain regions. For example, in the brain, it is assumed that the majority of neurons are cerebellum granule cells [48]. In [31], the number of synaptic connections to granule neurons is estimated to an average of only 4 connections, matching those observed anatomically. On the other hand, Purkinje neurons can have up to 200, 000 synapses on only one dendrite in the human brain [48]. The approximate number of synapses in the cortex is 0.6 · 10^14^ [10]. Assuming that the volume of the cortex represents around 80% of the volume of the brain, the number of synapses in the brain is of order 10^14^. Considering that the number of neurons in the human brain is of order 10^11^ [21], we find that the average number of synapses is about 1,000 synapses per neuron^3^. In the present work, the parameter *d*, the average number of pre / post synaptic neurons, is therefore evolving between 250 and 1000.

Finally, an action potential arriving on one pre-synaptic neuron produces an Excitatory Post-Synaptic Potential (EPSP), or an Inhibitory PostSynaptic Potential (IPSP), in the postsynaptic neuron. The duration of these postsynaptic potentials is about *τ* = 20*ms* [48] (see Section 2.1 for the link with the support of the interaction functions).

Therefore the parameters that we used in the simulation are indicated in Table 1.

**Table 1:** Parameters used in the simulation

|  |  |
| --- | --- |
| Simulation duration | $T = 5s$ |
| Mean firing rate | $\bar{m} = 0.3Hz$ |
| Number of neurons simulated | $M \in \{10^5, 10^6, 10^7, 10^8\}$ |
| Synaptic connection in average | $d \in \{250, 500, 1000\}$ |
| Postsynaptic duration | $\tau = 0.02s$ |

### 2.6 Software and hardware configurations

The simulations have been run on a Symmetric shared Memory multiProcessor (SMP) computer, called IRENE, equipped with Intel CascadeLake@2.6GHz processors^4^. This kind of computer is used here to have access to larger memory capacities. At computational level, only one PU (*i.e*. a single core running a single thread) was used for the sequential simulations. The implementation of the algorithm is written in C++ (2011) programming language and compiled using g++ 9.3.0.

### 2.7 Firing rate at network level

As detailed above, the firing rates in the brain are highly inhomogeneous. Therefore, to be more realistic, it was important to simulate this heterogeneity. Also this heterogeneity is advantageous to ATiTA(P) since the whole algorithm does not spend time on almost silent neurons to concentrate all the single-thread computations of the single PU on the most active neurons. Therefore, before looking at execution time and memory imprint we wanted to check this heterogeneity.

Table 2 presents classical elementary statistics on the simulated firing rates, whereas Figure 4 presents the corresponding densities. As one can see in Section 3, the system is initialized with a lot of neurons whose spontaneous spiking activity is null (see (1), with *ν_i_* = 0). The system needs to warm up to have almost all neurons spiking. This explains why the density at *T* = 5s is still rippled whereas, at *T* = 50s, it looks much smoother. As detailed in Section 3, the parameters of the Hawkes model (in particular the spontaneous spiking activity) have been fixed to achieve a certain stationary distribution of the firing rates (with mean 0.3Hz), which is heavy tailed to achieve records as large as 50 Hz. This is close to the distribution at *T* = 50 s. However these simulations show that, even if at *T* = 5s the system is not warmed up yet with a lot of non spiking neurons, one can still achieve the desired average firing rate and heterogeneity (extremal values) and that this does not vary a lot with *T* (see Table 2). Note that the density plots are roughly the same for all configurations: with ripples at *T* = 5s and smooth curves at *T* = 50s.

**Figure 4:**
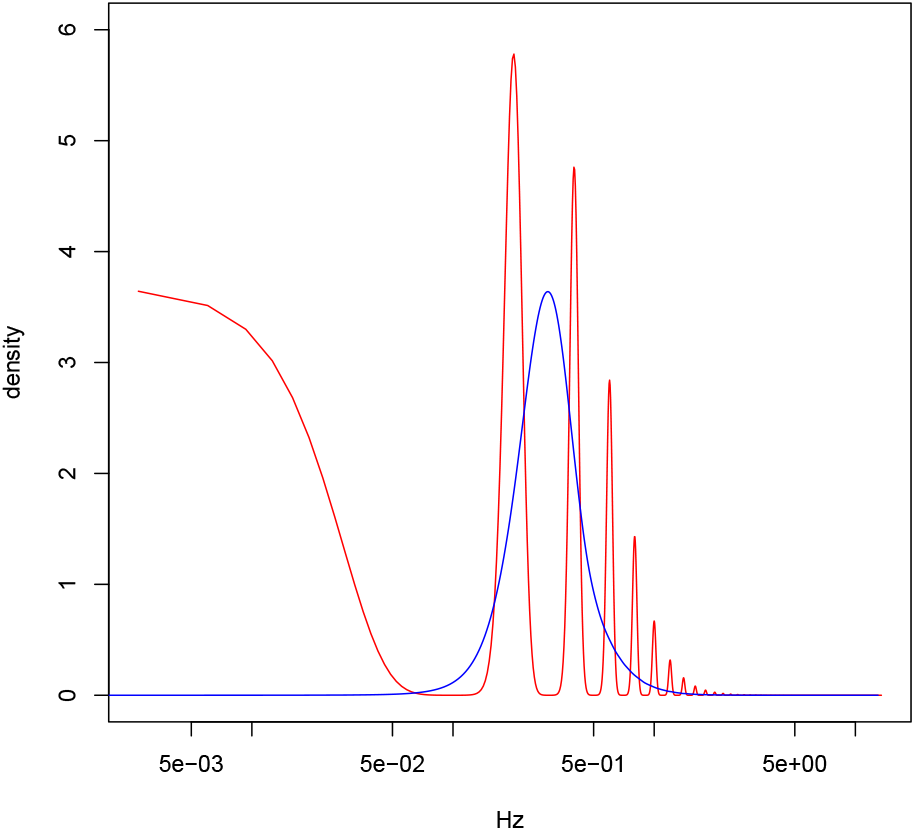
Densities (on a logarithmic scale) of the simulated firing rates in the network with *M* = 10^6^ neurons and *d* = 1000 post-synaptic connections in average. In red, for *T* = 5s and in blue for *T* = 50s. These densities are obtained with a Gaussian kernel estimator with bandwidth 0.02 Hz.

**Table 2:** Firing rates elementary statistics (average, minimum, maximum and standard deviation) obtained by simulation for different sizes of neural networks and different numbers of synaptic connections and *T* = 5s. The number between parentheses displays the results at *T* = 50s. We were able to run these long simulations of 50s only for the less demanding set of parameters (typically not 10^7^ neurons and 1000 synapses).

| M | d | Average freq. | Freq. min. | Freq. max. | Freq. std. | Percentage of non spiking neuron |
| --- | --- | --- | --- | --- | --- | --- |
| 1e5 | 250 | 0.279 (0.279) | 0 (0) | 14 (13.94) | 0.315 (0.222) | 31.2 (0.01) |
| 1e5 | 500 | 0.334 (0.333) | 0 (0.02) | 6.6 (5.76) | 0.328 (0.218) | 23.5 (0) |
| 1e5 | 1000 | 0.399 (0.398) | 0 (0.04) | 10.4 (11.64) | 0.345 (0.220) | 16.9 (0) |
| 1e6 | 250 | 0.267 (0.267) | 0 (0) | 19.2 (20.84) | 0.308 (0.217) | 33 (0.01) |
| 1e6 | 500 | 0.322 (0.324) | 0 (0) | 38.4 (39.38) | 0.329 (0.225) | 25.1 (0.00) |
| 1e6 | 1000 | 0.383 (0.387) | 0 (0.02) | 13.4 (12.9) | 0.344 (0.223) | 18.5 (0) |
| 5e6 | 250 | 0.26 (0.261) | 0 (0) | 27.4 (28.5) | 0.307 (0.217) | 34.1 (0.02) |
| 5e6 | 500 | 0.315 (0.316) | 0 (0) | 34.2 (34.1) | 0.324 (0.220) | 26 (0.00) |
| 5e6 | 1000 | 0.377 | 0 | 19.8 | 0.342 | 19 |
| 1e7 | 250 | 0.258 (0.259) | 0 (0) | 23.2 (21.62) | 0.306 (0.217) | 34.4 (0.02) |
| 1e7 | 500 | 0.311 | 0 | 21.6 | 0.322 | 26.4 |
| 1e7 | 1000 | 0.374 | 0 | 21.8 | 0.342 | 19.3 |
| 5e7 | 250 | 0.253 | 0 | 38.2 | 0.304 | 35.4 |
| 5e7 | 500 | 0.305 | 0 | 50.6 | 0.321 | 27.2 |
| 1e8 | 250 | 0.251 | 0 | 46.2 | 0.303 | 35.7 |

### 2.8 Execution times and memory usage

Equation (2) has a leading term which is mainly 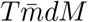 log(M), as soon as 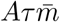 is small, which is the case in realistic simulations.

The simulation execution times are presented in Figure 5a for different *M* (number of neurons) and different *d* (average number of pre/post synaptic connections). The experimental execution times obtained are in agreement with (2): the curves (in the log scale) are almost linear, with slopes around 1.1, which corresponds to the theoretical growth in *dM* log(*M*) with respect to the number of neurons. Also since *d*, the number of post-synaptic connections, acts like a multiplicative factor, this explains the parallel equispaced lines in the log-scale. Finally, Figure 5c shows that the ratio of execution times is roughly around 10, which is in accordance with the linear dependency in *T*.

**Figure 5:**
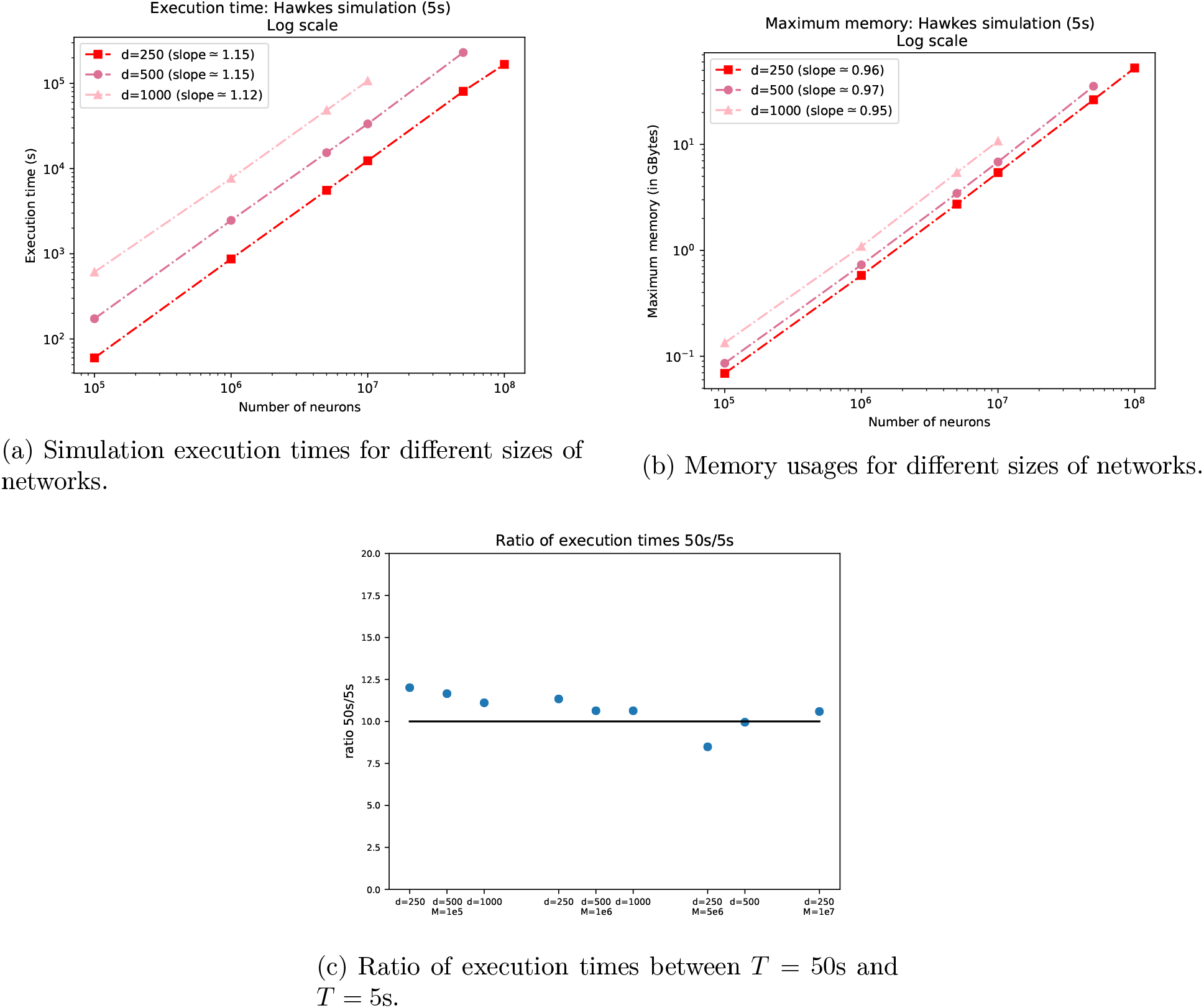
Simulation execution times (5a) and memory usage (5b) of ATiTAP for different sizes of networks and different simulation durations.

The total amount of used memory is displayed in Figure 5b. They are also in agreement with ATiTAP theoretical memory complexity, computed in Equation (4): it is almost linear in *M*, with slopes around 1, whatever *d*.

Note that a mouse cortex or a human hippocampus have roughly *M* = 10^7^ neurons [54]. Figure 5 says that in this case, with *d* = 1000, we need a few hours and about 10 GigaBytes of memory to simulate it. Because of the configuration we used, Figure 5 therefore shows that this could have been achieved on a simple desktop computer.

## 3 Material and Method

### 3.1 Details on the model

For a set of *M* neurons, we first design the graph of interactions by saying that neuron *j* influences neuron *i* if a Bernoulli variable *Z_j→i_* of parameter *p* is non zero. The resulting network is an Erdös-Rényii graph. Once the network is fixed, we design the spike apparition based on a Hawkes process (see (1), with Φ(*x*) = *x*).

One of the advantage of (linear) Hawkes processes is that one can evaluate the firing rate distribution and that this evaluation only depends on the integral of the interaction function. In this sense, having piecewise constant functions, or shifted piecewise constant functions to take into account synaptic delays, or even more intricate (like exponential etc.) interaction functions, will absolutely not change the evaluations that are made as long as the integral is known (see Figure 1).

We are interested in a particular case of the Hawkes process where all the interaction functions are always the same when they are non null. More precisely, we set the interaction function

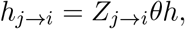

where *h* is a fixed positive interaction function of integral 1 and *θ* is a tuning parameter that we need to calibrate to avoid explosion of the process. We also set *h_i→i_* = 0 (no self interaction). We take 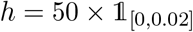, so that *h* is of integral 1 and non zero *h_j→i_* are of integral θ.

Let us denote 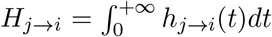 and *H* = (*H_j→i_*)_*i,j*=1,…, *M*_ the corresponding matrix (line *i* corresponds to a postsynaptic neuron, column *j* to a presynaptic neuron).

Hawkes processes may explode that is, it produces an exponentially increasing number of points per unit of time (see [8]), but this does not happen if the spectral radius of *H* is strictly smaller than 1 [20]. In this case, a stationary version exists and the corresponding vector of mean firing rates *m* = (*m_i_*)_*i*=1,…, *M*_ is given by

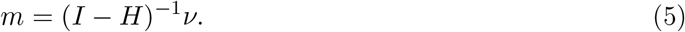

Note also that if we start the simulation without points before 0 in this case, the process is not stricto sensu stationary but it converges to an equilibrium given by the stationary state and that the number of points that are produced in this case is always smaller than the stationary version.

Before running the simulation, we want to calibrate parameters so that (i) we avoid explosion and (ii) we reach a certain realistic heterogeneous distribution of the firing rates (vector *m*), that is an average around 0.3 Hz and records around 50 Hz. Both of these calibrations can be done mathematically beforehand in the Hawkes model: we can guarantee the behavior of the whole system even before performing the simulation, whereas this might be much more intricate for other models such as LIF.

### 3.2 Choice of *θ* or how to avoid explosion

Note that 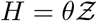, with 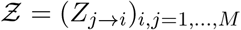. So if we can compute the largest eigenvalue of 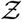 or an upper bound, we can decide how to choose *θ*.

We can use Gershgorin circles [53] to say that any complex eigenvalue *λ* of 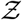 satisfies (because the diagonal is null),

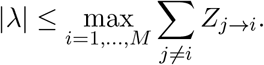

Therefore the spectral radius is upper bounded by max_*i*=1,…,*M*_ *B_i_*, where *B_i_* = ∑_*j*≠*i*_ *Z_j→i_*. This random quantity can be computed for small networks but it is clearly too intensive in our setting: indeed, with the procedural connectivity implementation, it is always easy to access the children *ℓ* of a given *i*, *i.e*. such that *i* → *ℓ* is in the graph, but we need to look at all the neurons in the graphs to find out the set of parents *j* of *i*, *i.e*. such that *j* → *i* is in the graph. However, probabilistic estimates might be computed mathematically. Indeed *B_i_* is a sum of i.i.d. Bernoulli variables. So by Bernstein’s inequality [1], for all positive *x*,

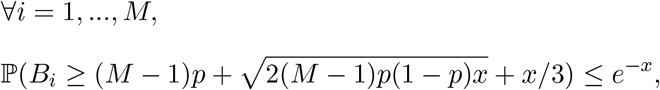

and, by union bound,

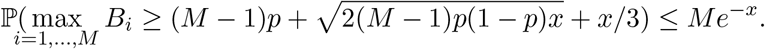

Therefore for a fixed level *α* (say 1%), and with *x* = log(*M*) + log(1/*α*), we obtain that with probability larger than 1 – *α*, the spectral radius of 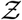 is upper bounded

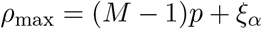

with

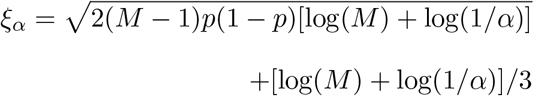

Note that *ρ_max_* is roughly (*M* – 1)*p*, which is the largest eigenvalue of 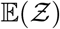. Finally if we take *θ* < 1/*ρ*_max_, the process will not explode with probability larger than 1 – *α*. In practice, to ensure a strong enough interaction, we take 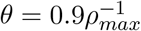.

#### Choice of *ν_i_* or how to constraint the distribution of the firing rates

The first step consists in deciding for a target distribution for the *m_i_*’s. We have chosen to pick the *m_i_*’s independently as 0.1*X* where *X* is the absolute value of a student variable with mean 3 and 4 degrees of freedom. The choice of the student variable was driven by the wish of having a moderate heavy tail, which will ensure records around 50 Hz and a mean around 0.3Hz.

The problem is that the *m_i_*’s are not parameters of the model, so we need to tune the *ν_i_*’s to get such *m_i_*’s. Note that by inverting (5), we get that

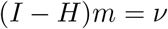

(see also [43]) that is for all *i*

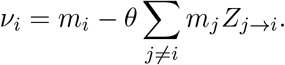

This means that the spontaneous rate that we need to put is the mean firing rate *m_i_* minus what can be explained with the parents of *i*.

So in theory, the Hawkes model is very easy to tune for prescribed firing rates since there is a linear relationship between both. However, and for the same reasons as before, it might be too computationally intensive to compute this explicitly.

One possible way is to again use concentration inequalities, but this time on ∑_*j≠i*_ *m_j_ Z_j→i_* and not on *B_i_*. However a simpler trick works well (as seen in Figure 4).

Indeed ∑_*j≠i*_ *m_j_Z_j→i_* is a sum of about (*M* — 1)*p* ≃ 1000 i.i.d variables with mean 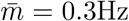. Hence it should be close to 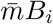. With the previous computations, we know already that *v_i_* should therefore be larger than 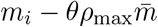.

Therefore, with the previous choice of 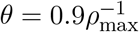, we take the *v_i_*’s as follows:

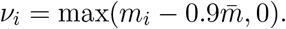

Note that *ν_i_* remains positive or null, which guarantees that the Hawkes process stays linear. However, this also means that a non negligible portion of the neurons start with a null spontaneous firing rate, which explains the ripples of Figure 4.

With this choice, we cannot hope to have exactly the same distribution as the desired *m_i_*’s, but it conserves the same heavy tail and roughly the same mean firing rate, as one can see on Table 2.

## 4 Discussion

The ATiTAP algorithm is based on Activity Tracking with Time Asynchrony and Procedural connectivity and is derived from another algorithm, ATiTA [32] which has a much higher memory cost. The aim of this study is a proof of concept that simulations of huge neuronal networks on single PU are possible and that this may open the way for a new point of view on neuronal simulations.

More specifically, we have been able to theoretically compute ATiTA(P) computational cost and memory burden (see Sections 2.3 and 2.4) and to perform simulations in a range of parameters that are of the order of magnitude of human brain areas or small mammalian whole brains (see Section 2.5). Moreover we have been able to reproduce a large heterogeneity in the firing rates (see Section 2.7) and to prove that on a desktop computer we can easily reach 10^7^ neurons, that is the size a human hippocampus [54] or of a mouse.

Comparison with existing algorithms is tricky. First because the models are not completely equivalent (for instance Integrate-and-Fire in [19, 26] versus Hawkes for ATiTA(P)), second because the other algorithms use supercomputers or GPUs with massive parallel computations (*cf*. Table 3), thirdly because depending on the model, it is more or less difficult to compute theoretical memory and computational costs (see for instance [27, 24]).

**Table 3:** Comparison of execution times and total memory cost for each simulator and comparable network sizes for 1s of biological time. For ATiTAP (see Section 2 and Figure 5) the case *d* = 1000, *M* = 10^6^ corresponds to a total number of synapses of 10^9^. Note that in **NEST1** and **NEST2**, synapses are plastic, that is, a certain form of update of the synaptic weights is embedded in the code and the whole network should be stored (this is also the case for ATiTA). The last two rows correspond to what might have been the execution times and memory cost if we implemented ATiTA(P) but with *d* = 10^4^, that is 10^10^ synapses. To understand the theoretical scaling for ATiTAP and ATiTA with 10^10^ synapses, we apply for the computational cost, the theoretical multiplicative factor given by the ratio of (2) between *d* = 10^4^ and *d* = 10^3^. The same computation is done for the memory cost with (4) in the case where *ρ* and *η* are neglected and where *ω* = *ϵ*, that is same accuracy for a spiking time and an index in the network. **NEST1** and **NEST2** used 1 MPI process per node. For the number of threads of **NEST1** and **NEST2**, we thus multiplied the number of MPI processes that have been used by the number of used threads per node. For the memory usage of **NEST1** and **NEST2**, we multiplied the code memory usage per node, 2 GiBytes and 12 GiBytes respectively, by the number of nodes used for the simulations. For **GeNN**, the indicated number of threads is the number of CUDA cores since this simulation is done on GPU cards. When several implementations were possible, we took each time the one with the smallest execution time.

| Simulator | Machine | Size order | Memory | Execution time | Nb. Threads |
| --- | --- | --- | --- | --- | --- |
| **NEST1** [47] | JUQUEEN | $10^6$ neurons<br>$10^{10}$ synapses | $2 \cdot 10^{12}$ Bytes | 17min | $1024 \cdot 64 = 65536$ |
| **NEST2** [24] | JUQUEEN | $10^6$ neurons<br>$10^{10}$ plastic synapses | $10^{14}$ Bytes | 16s | $8192 \cdot 8 = 65536$ |
| **GeNN** [26] | GPU | $10^6$ neurons<br>$10^{10}$ synapses | $1.2 \cdot 10^{10}$ Bytes | 14.4min | 4608 |
| **ATiTAP** | IRENE | $10^6$ neurons<br>$10^9$ synapses | $10^9$ Bytes | 25min | 1 |
| **ATiTAP** (Th.) | IRENE | $10^6$ neurons<br>$10^{10}$ synapses | $8.7 \cdot 10^9$ Bytes | $36 \cdot 25 \text{mn} = 15\text{h}$ | 1 |
| **ATiTA** (Th.) | IRENE | $10^6$ neurons<br>$10^{10}$ synapses | $2.8 \cdot 10^{12}$ Bytes | $36 \cdot 25 \text{mn} = 15\text{h}$ | 1 |

In Section 2.1, we have shown the similarities and dissimilarities between Hawkes processes and Integrate-and-Fire. We believe that it is a matter of taste for preferring one to the other: Hawkes processes have been shown to fit real data (see [44, 41]), Integrate-and-Fire models have a nice voltage interpretation. But there are links between both, especially because the intensity of the process can be thought as a function of the voltage.

Besides the fact that Hawkes processes allow us to compute theoretical complexities, we believe that one of the main advantage of Hawkes processes is that they satisfy time asynchrony, which allows us to spend computational time only on the neurons that are very active. Let us quantify this more specifically. In Section 2.2, we have described in details the benchmark hybrid algorithm [19] versus ATiTA(P). The main burden with parallelization, even in the hybrid algorithm, is that PUs have to wait for each other every discretization step *T_com_*. The nice idea of the hybrid algorithm is that this step is not the thinnest possible but of intermediate size, so that this synchronization does not happen often (*T_com_* = 0.001s in their case). Still, at every discretization step, the whole system has to check the state of all neurons, that is a total of *MT/T_com_* operations just for the synchronization. For a classic network of *M* = 10^6^ neurons and 1s of simulation, we end up with 10^9^ operations, only for synchronization. To give an order of magnitude, ATiTA and ATiTAP have a complexity cost of 6 · 10^9^ thanks to (2) for 1000 post-synaptic neurons in average, for the full simulation.

Table 3 compares both execution times and memory imprints of our simulation with the latest simulations of large-scale spiking neural networks in the literature^5^. Because the models are very close dynamically and structurally, it might be a good way to roughly compare the memory and computational performances of corresponding simulators and hardware solutions. Structurally, networks of similar size and connectivity are compared. Dynamically, simulations in [26, 47, 24] use LIF models that are dynamically very close to the Hawkes model used here (see Section 2.1). The simulations we found in the literature have about *d* = 10^4^ synapses and we used the theoretical complexities to give an idea of how ATiTA(P) would scale. However we need to underline that comparing completely these models is very tricky. First of all, they have very different firing rates (up to 14.6 Hz in some of the models used in [47]). A first look at the main term 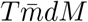 log(M) in the theoretical computational complexity might let us think that we need to multiply by about 50 these execution times. However, we think this crude estimate might be biased, especially because the models in [26, 47] are representing with a lot of details the visual cortex with different layers, each layer having a different connectivity and number of neurons. For these highly heterogeneous models, a mean firing rate and mean number of synapses might not represent to a full extent what ATiTA(P) can do and we refer the interested reader to [32] for a more detailed formula in the case of structured networks.

In any case, despite the difficulty of the exact comparison, we see that ATiTA(P) as GeNN, the two algorithms implementing procedural connectivity, have a comparable and significant decrease in memory imprints with respect to the other algorithms and that ATiTA (that is, the algorithm that stores the connectivity graph), would have had a memory imprint comparable to NEST1 and smaller than NEST2. Note that storing all the synaptic weights is a bottleneck that cannot be easily passed as soon as one wants to simulate synaptic plasticity, which is not the purpose of the present work.

In terms of execution times, ATiTA(P) is clearly the slowest with a factor 3 · 10^3^ with respect to the fastest one (NEST2)^6^, but ATiTA(P) only uses one thread with respect to the 6 · 10^4^ threads used in NEST2. This has to be put in perspective with respect to the purpose of simulations. Indeed, especially for random models (but this can also hold as soon as we want to test several set of parameters), the purpose of simulations is not to make just one simulation but to do replicas. In this sense, it is important to think that if we use the totality of the threads to perform one simulation, doing replicas can only be done in a serial way. In this sense, ATiTA(P) allows us to have one thread per replica and one can perform parallel simulations of the different replicas very easily. So even if the comparison in terms of execution times is purely qualitative, due to the heterogeneity of the models that have been simulated. Table 3 shows that roughly speaking, ATiTA(P) can simulate as many replicas as NEST2 in about the same amount of time: ATiTA(P) would use the 6 · 10^4^ threads to simulate the replicas, whereas NEST2 would use these threads to simulate faster one replica and then would implement sequentially the various replicas. Notice that the number of replicas can be limited by the memory capacity of the supercomputer used to run the replicas. Indeed, each core hosting the threads should have enough memory to actually host an ATiTAP simulation. Taking JUQUEEN supercomputer to run ATiTAP replicas, each node has 16 GiBytes of memory. This allows 16GB/8Threads = 2 GiBytes of available memory per running thread. An ATiTAP instance consuming 8.7 GiBytes of memory according to Table 3. Thus one ATiTAP instance actually consumes the memory resources of 8.7GiBytes/2GiBytes per core, *i.e*., 4 cores. This actually reduces the possible number of parallel replicas run by ATiTA(P) by a factor of 4: 65536/4=16384. The same could roughly apply to GeNN as well but notice that GPUs are designed to run one simulation in parallel, they are not adapted to run several simulations in parallel. For example, the Nvidia Titan RTX used by GeNN only has a total memory of 24 GiBytes allowing only running 2 replicas in parallel.

The present work is therefore a proof of concept that optimized sequential algorithms based on activity tracking and time-asynchrony might well offer another point of view on massive simulations of brain-scale neural networks.

There is still room for improvements. Among the several possible improvements, ATiTA(P) could be parallelized among the post-synaptic neurons (update and prediction of the next potential spikes). This is particularly important if we want to pass from *d* = 10^3^ to *d* = 10^4^ post-synaptic neurons in average (even if we showed that *d* = 10^4^ seems not to be relevant for realistic simulations). Next, new algorithms for Hawkes processes, based on Kalikow decomposition, have been proved to work even if the neuronal network is infinite [40]. This would theoretically decrease the overall complexity in terms of *d*. Finally, if we want to have synaptic plasticity, we are forced to work with ATiTA and varying non zero interaction functions that need to be stored. Once they are stored, the update of these functions seems to be as costly that the update of the intensities themselves (see for instance [25] where only local firing rates are needed). Once all these improvements are done, one might think to implement fully realistic models (such as the one of the visual cortex of [47]) with variants of ATiTA(P). This is definitely a source for future work, which might lead to simulations, whose execution time is smaller than biological time [28].

## Acknowledgements

We would like to thank the anonymous reviewers, whose remarks allowed to greatly improve the quality of this article.

This work is part of the project HyperBrain from Human Brain Project (HBP) EBRAINS EU initiative. The simulations were run on Fenix Infrastructure resources, which are partially funded from the European Union’s Horizon 2020 research and innovation program through the ICEI project under the grant agreement No. 800858. Our research was supported by the French government, through CNRS, the UCA^Jedi^ and 3IA Côte d’Azur Investissements d’Avenir managed by the National Research Agency (ANR-15-IDEX-01 and ANR-19-P3IA-0002), directly by the ANR project ChaMaNe (ANR-19-CE40-0024-02) and by the interdisciplinary Institute for Modeling in Neuroscience and Cognition (NeuroMod) of the Université Côte d’Azur.

## Footnotes

1 Note that by “continuous” we mean that it is possible to compute the spiking time up to the usual double numerical precision of 10^-15^. Indeed, as indicated in [19] this numerical precision can be decoupled from the global computation time step, which corresponds to the minimal synaptic delay *T_com_*. As shown in [19], for a certain accuracy, the hybrid simulation scheme with spikes in continuous time is much faster than a hybrid approach where the computation time step limits the accuracy.

2 This calculus can be found on AI impact project webpage: https://aiimpacts.org/rate-of-neuron-firing/ (lastly verified: 02/09/2021)

3 Calculus on AI impact project webpage: https://aiimpacts.org/scale-of-the-human-brain/ (lastly verified: 02/09/2021).

4 We used the partition v100l on Joliot-Curie supercomputer at TGCC as a Fenix Infrastructure resource. A dualsocket mother board contains the 2 CPUs, Intel CascadeLake@2.6GHz processors, each with 18 cores. Sequential simulations were run using only a single core. Each core has a memory of 10 GBytes, so the total amount of available memory is 360 GBytes.

5 Notice that in [24] two supercomputers, JUQUEEN and K, were used but we present here JUQUEEN results, which are the most comparable ones. In [26] different GPU graphics cards were used however we refer here to the best-result one, the Nvidia Titan RTX.

6 If we take into account the firing rate, we might want to multiply this by a factor 50, but as said before, this might not be realistic.

